# Sevoflurane inhibition of the developmentally expressed neuronal sodium channel Na_v_1.3

**DOI:** 10.1101/2025.04.10.648240

**Authors:** Jiaxin Xiang, Karl F. Herold, Jimcy Platholi, Hugh C. Hemmings

## Abstract

Neuronal voltage-gated sodium channels (Na_v_) are major targets for the neurophysiological actions of general anesthetics. In the adult brain, cell type-specific effects on synaptic transmission are attributed to the differential sensitivity to volatile anesthetics of specific Na_v_ subtypes preferentially expressed in mature neurons (Na_v_1.1, Na_v_1.2, Na_v_1.6). Compared to mature neurons, neurons in the developing CNS are more excitable. Since the subtype-selective effects of volatile anesthetics on Na_v_ during early development are unknown, we determined volatile anesthetic effects on Na^+^ currents mediated by Na_v_1.3, the principal Na_v_ subtype expressed in developing neurons. Sevoflurane at clinically relevant concentrations inhibited peak Na^+^ current (*I*_Na_) of human Na_v_1.3 heterologously expressed in HEK293T cells in a voltage-dependent manner, induced a −6.1 mV hyperpolarizing shift in the voltage dependence of steady-state inactivation, and slowed recovery from fast inactivation. Na_v_1.3-mediated Na^+^ currents also exhibited distinct activation properties associated with neuronal hyperexcitability, including prominent persistent currents and ramp currents, both of which were significantly reduced by sevoflurane. The major neuronal subtype Na_v_1.2 showed a more hyperpolarized voltage dependence of steady-state inactivation than Na_v_1.3. Consistent with its lower propensity for sustained repetitive firing, Na_v_1.2 exhibited minimal persistent and ramp currents, and these were unaffected by sevoflurane. These findings identify subtype-specific effects of the volatile anesthetic sevoflurane on neuronal Na_v_ subtypes and suggest a mechanistic basis for increased anesthetic sensitivity in early neuronal differentiation and maturation.

## Introduction

Voltage-gated Na^+^ channels (Na_v_), which regulate neuronal excitability^1-4^, action potential (AP)- driven Ca^2+^ influx, and Ca^2+-^dependent neurotransmitter release^3,5-8^, are important targets for general anesthetics ^9-12^. Upon neuronal depolarization, Na_v_ rapidly activate then inactivate forming an AP, followed by a return to resting membrane potential^13^. Volatile anesthetics (VA) directly inhibit heterologously expressed Na_v_ ^11^ with depression of AP amplitude and reduce nerve terminal depolarization in neuronal preparations^14-17^, including dissociated neurons from the hippocampus^18^, a major site of anesthetic actions ^19^.

Four major Na_v_ subtypes (Na_v_1.1, Na_v_1.2, Na_v_1.3, Na_v_1.6) are expressed in the mammalian brain^20,21^, with Na_v_1.3 preferentially expressed during early development ^22,23^. In the hippocampus, variable expression of Na_v_ subtypes in specific neuronal populations influences cellular and network excitability with differential presynaptic and postsynaptic localization^8^. Mature inhibitory hippocampal interneurons are enriched in Na_v_1.1 compared to excitatory pyramidal neurons that express relatively more Na_v_1.2 and Na_v_1.6^24-26^. These Na_v_ subtypes are inhibited by VAs in two ways: 1) stabilization of the inactivated state resulting in a shift of steady-state inactivation to more negative membrane potentials, reducing channel availability and slowing recovery from fast-inactivation, and 2) interaction with the open and/or resting state to produce tonic block^9,27,28^. Voltage- and frequency-dependent block of Na_v_ ^17,28,29^ reduces AP amplitude^15,30^ and neurotransmitter release^31,15^ with Na_v_ subtype and neurotransmitter selectivity. For example, isoflurane decreases glutamate release more potently than GABA release from dissociated hippocampal neurons^32^, which is likely mediated through greater inhibition of Na_v_1.2 and Na_v_1.6 in glutamatergic boutons compared with lesser inhibition of Na_v_1.1 in GABAergic boutons^18,33^.

Heterogeneity in Na_v_ expression also occurs during neuronal differentiation and development. Expression of Na_v_ subtypes with distinct activation kinetics renders developing neurons hyperexcitable compared to mature neurons ^22^. Compared to other subtypes, Na_v_1.3 recovers from inactivation four-fold faster and has slower closed-state inactivation, reducing the threshold for activation in neurons that express Na_v_1.3, leading to sustained repetition firing^34-37^, even with unchanged or absent stimuli. Slower inactivation or resurgent Na^+^ currents enhance repetitive firing and modulate overall neuronal excitability as opposed to AP initiation and propagation^38,39^. Although persistent Na^+^ currents in acute hippocampal slices are inhibited by isoflurane^40^, subtype-selective effects of anesthetic sensitivity on developmentally expressed Na_v_ have not been investigated. We examined the effects of sevoflurane on the function of Na_v_1.3, a major neuronal subtype expressed in the developing mammalian brain. Comparative studies with Na_v_1.2 provide a neurophysiological basis for developmental stage-specific impact on neuronal excitability.

## Materials and Methods

### Materials

Sevoflurane was obtained from Henry Schein Medical (Melville, NY). HEK293T cells were from ATCC (Manassas, VA). All other chemicals were from Sigma-Aldrich (St. Louis, MO).

### cDNA Constructs

Human Na_v_1.2-pIR-CMV-IRES-mScarlet (Accession number NM_021007) and human Na_v_1.3- pIR (Accession number NM_001081676) cDNA were kindly provided by Professor A.L. George Jr. (Northwestern University, Evanston, IL) via AddGene (Watertown, MA) and Professor J. A. Kearney (Northwestern University, Evanston, IL), respectively.

### Cell culture and electrophysiological recordings of Na^+^ currents

Human HEK293T cells (#CRL-3216, ATCC, Manassas, VA) were seeded into 35-mm dishes in Dulbecco’s Modified Eagle′s Medium (Sigma-Aldrich) supplemented with 10% (v/v) heat-inactivated fetal bovine serum, 100 U/ml penicillin, and 100 mg/ml streptomycin and incubated in a humidified atmosphere of 5% CO_2_ and 95% air at 37°C. HEK293T cells were cotransfected with cDNA for hNa_v_1.2 or hNa_v_1.3 (3-4 μg) and human auxiliary hβ1 subunit (0.8-1 μg) using Lipofectamine LTX (Invitrogen, Carlsbad, CA). Additionally, hNa_v_1.3 was cotransfected with 0.8 μg pEGFP-N1 (Clontech, Mountain View, CA) as a reporter to allow identification of EGFP-transfected cells by fluorescence microscopy. Cells were re-seeded at lower density onto 12- mm glass coverslips 20-24 h post-transfection, and electrophysiological studies were conducted at least 3 h later to allow cells to adhere.

For electrophysiological recordings, coverslips were transferred into a small volume laminar-flow perfusion chamber (Warner Instruments, Hamden, CT) and continuously perfused at ∼2 ml/min with extracellular solution containing (in mM): 130 mM NaCl, 10 mM HEPES, 3.25 mM KCl, 2 mM MgCl_2_, 2 mM CaCl_2_, 0.1 mM CdCl_2_, and 20 mM TEA-Cl, adjusted to pH 7.4 (with NaOH) and to 310 mOsm/kg. Na^+^ currents were recorded in voltage-clamp mode at room temperature (23– 24°C) using an Axopatch 200B patch-clamp amplifier (Molecular Devices, San Jose, CA) with a 5 kHz low-pass filter at a sampling rate of 20 kHz. Recording pipettes were pulled from borosilicate glass capillaries (Sutter Instruments, Novato, CA) on a P-1000 horizontal puller (Sutter Instruments) and fire-polished before use. Pipette resistance was ∼1.5-2.5 MΩ when filled with internal solution containing (in mM): 120 mM CsF, 10 mM NaCl, 10 mM HEPES, 10 mM EGTA, 10 mM TEA-Cl, 1 mM CaCl_2_, and 1 mM MgCl_2_ adjusted to pH 7.3 (with CsOH) and to 312 mOsm/kg (with sucrose). Access resistance (∼2–4 MΩ) was reduced using 75–85% series resistance correction. To minimize space-clamp and series resistance errors, only cells expressing 1–8 nA peak current were analyzed. Initial whole cell seal resistance was 1–3 GΩ, and recordings were discarded if resistance dropped below 1 GΩ. Liquid–junction potentials were not corrected. Capacitive current transients were electronically canceled with the internal amplifier circuitry, and leak currents were digitally subtracted using the P/4 protocol^41^ Recordings began 5 minutes after attaining the whole-cell patch configuration to allow equilibration of pipette solution and cytosol. Stimulation protocols were applied in control external solution, and again after a 3- minute superperfusion with external solution containing sevoflurane.

### Sevoflurane superfusion

A saturated stock solution of sevoflurane in external solution (∼5.2 mM at 23 °C) was diluted to the desired final concentration in a gas-tight glass syringe. Sevoflurane solutions were superfused using a pressure-driven microperfusion system (ALA Scientific, Westbury, NY) with a 200 μm diameter perfusion pipette tip positioned ∼200 μm from the recorded cell. Sevoflurane concentrations sampled at the perfusion pipette tip were determined by gas chromatography using a Shimadzu GC-2010 Plus gas chromatograph (Shimadzu, Tokyo, Japan) after extraction into octane ^42^. An aqueous sevoflurane solution of 0.28 mM was used as the predicted minimum alveolar concentration (MAC; equivalent to the EC_50_ for immobilization) in humans after temperature adjustment to 24°C^30,43^.

### Stimulation protocols and data analysis

The holding potential (*V*_h_) was –80 mV unless otherwise stated. Voltage-dependent inhibition of peak inward Na^+^ current (*I*_Na_) was analyzed using a 20 ms test pulse to 0 mV preceded by a 100 ms prepulse alternating between either –120 mV (*V*_0_) or the voltage of half-maximal inactivation (*V*_½-inact_ = –47±5 mV) applied every 5 s^10,44,45^. *V*_½-inact_ was determined for each individual cell using the double-pulse protocol for steady-state inactivation described below. For voltage-dependence of activation, conductance (*G*) values were derived from the current-voltage (I-V) relationship using:

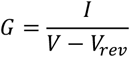

where *I* is the peak sodium current (*I*_Na_) at a given command voltage (*V*), and *V*_rev_ is the measured Na^+^ reversal potential. Conductance values were normalized to maximal conductance (*G*/*G*_max_), and the activation curve was fitted using a Boltzmann function:

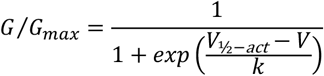

where *V*_½act_ is the voltage at which activation is half-maximal, and *k* is the slope factor describing the voltage sensitivity of activation.

Steady-state fast inactivation (*h*_∞_), was measured using a double-pulse protocol with a 100 ms prepulse ranging from –110 to +30 mV in 10 mV steps, followed by a 20 ms test pulse to 0 mV. Peak currents during the test pulse were normalized to the maximal current (*I*_Na_/*I*_Na_max, where *I*_Na_max is the maximal current that is elicited at the test potential), plotted against the prepulse potential (*V*_m_), and fitted with a Boltzmann function:

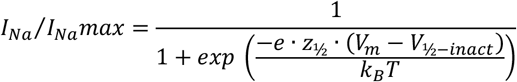

where *z*_½_ and *V*_½-inact_ denote the apparent gating valence and potential for half-maximal inactivation, respectively.

Recovery from fast inactivation was tested from a holding potential of –130 mV using a two-pulse protocol with two identical 10 ms pulses to 0 mV separated by an inter-pulse interval with increasing durations from 1–200 ms. Current amplitudes were normalized (Pulse_2_/Pulse_1_), plotted against inter-pulse interval. The recovery time course was best fitted with a bi-exponential equation to obtain the recovery time constants τ:

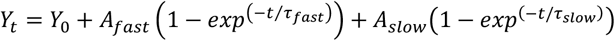

where *Y*_t_ denotes the normalized current at time *t, Y*_0_ denotes the normalized current at zero time, *A*_fast_ and *A*_slow_ represent the amplitudes of the fast and slow components, *t* is the inter-pulse interval, and τ_fast_ and τ_slow_ denote the fast and slow recovery time constants, respectively.

Statistical significance was assessed by two-tailed paired Student *t* test. *P* < 0.05 was considered statistically significant. Programs used for data acquisition and analysis were pClamp 10 (Molecular Devices), Excel (Microsoft, Redmond, WA), and Prism 10 (GraphPad Software Inc., La Jolla, CA). Values are reported as mean ± SD unless otherwise stated.

## Results

### Effects of sevoflurane on peak Na^+^current

The effects of sevoflurane on Na_v_1.3 currents were tested using a periodic stimulation protocol that allowed us to determine voltage-dependent drug effects^10,44,45^. To assess peak Na^+^ current (*I*_Na_) inhibition, a depolarizing pulse to 0 mV was applied every 5 s, following a 100 ms prepulse to either *V*_0_ (–120 mV), where most channels remained in an available resting state, or to a potential at which ∼50% of channels were in an inactivated state *V*_½-inact_ (–47±5 mV). Figure 1B shows the time course of *I*_Na_ inhibition during sevoflurane wash-in. The slow onset likely reflects drug partitioning into intracellular compartments rather than an intrinsically slow binding rate to the channel^46^. Results are expressed as the ratio of peak *I*_Na_ in the presence of sevoflurane to control *I*_Na_ (Fig. 1, Supplemental Table S1). Sevoflurane significantly inhibited peak *I*_Na_ following a prepulse to *V*_½-inact_ in a concentration-dependent manner.

**Fig. 1.**
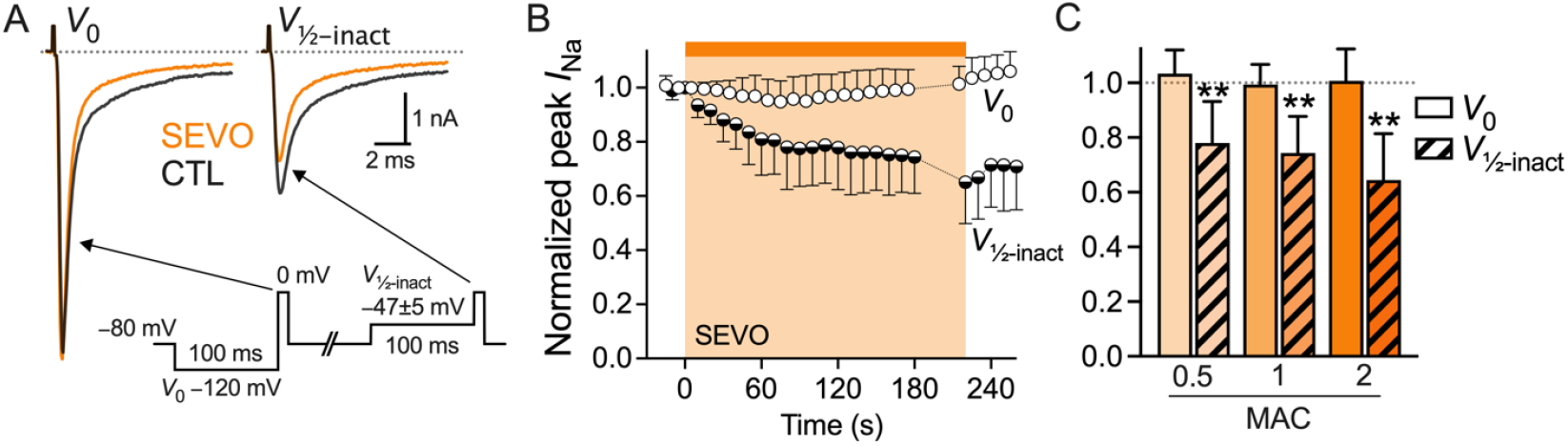
Inhibition of peak Na_v_1.3 current (*I*_Na_) by sevoflurane. Sevoflurane inhibited peak Na^+^ current (*I*_Na_) in a voltage-dependent manner at concentrations equivalent to 0.5 MAC, 1 MAC, or 2 MAC (0.14 mM, 0.28 mM, or 0.57 mM). (*A*) Macroscopic Na^+^ currents from a transfected HEK293T cell expressing Na_v_1.3 were evoked using a periodic alternating stimulation protocol (see *inset*) in the absence (CTL, *black traces*) or presence (SEVO, *orange traces*) of 1 MAC sevoflurane (0.28 mM). Sevoflurane did not inhibit peak *I*_Na_ when the test pulse followed a prepulse to *V*_0_, a potential at which all channels were in the resting state. However, sevoflurane significantly inhibited peak *I*_Na_ with a prepulse to *V*_½-inact_, a potential at which half of the channels were in a fast-inactivated state. (*B*) The time course of the effects of wash–in and wash–out of 1 MAC sevoflurane. (*C*) Concentration-dependent inhibition by sevoflurane at 0.5, 1, and 2 MAC (for values, see Supplemental Table S1). All data are mean±SD, n = 6-7, **P < 0.01 *vs*. control by paired two-tailed Student *t*-test.

### Effects of sevoflurane on activation and inactivation

Sevoflurane caused a small but significant hyperpolarizing shift in the voltage dependence of channel activation, indicating that channels opened at slightly more negative potentials (Fig. 2A,B). This suggests that sevoflurane facilitates activation by lowering the voltage threshold for channel opening. Steady-state fast inactivation, also referred to as channel availability (or *h*_∞_ from the Hodgkin-Huxley model), was tested using a double-pulse protocol (Fig 2A, *inset*). Sevoflurane significantly shifted the voltage dependence of steady-state inactivation by −6.1±2.7 mV toward more hyperpolarized potentials (Fig. 2C, Supplemental Table S1). This indicates that a greater fraction of channels either transitioned into and/or remained in the inactivated state at any given potential, reducing the pool of available channels for subsequent activation. The shift in *V*_½-inact_ was concentration-dependent, with the largest effect observed at 2 MAC (Fig. 2D). These findings are consistent with sevoflurane stabilization of the inactivated state, thereby limiting Na_v_ availability, while also slightly modulating activation kinetics.

**Fig. 2.**
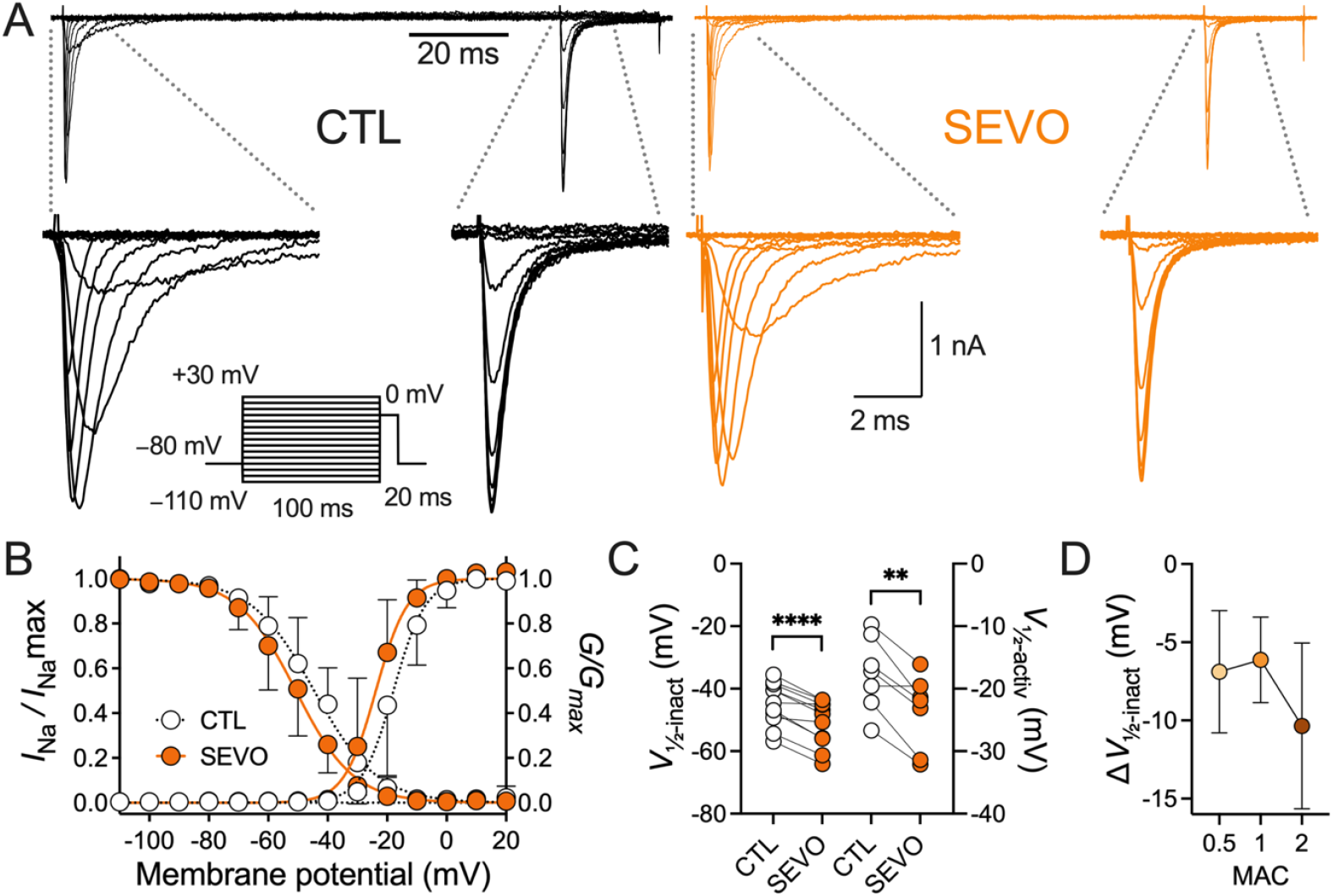
Effects of sevoflurane on Na_v_1.3 activation and inactivation. (*A*) Representative families of whole- cell inward *I*_Na_ evoked by depolarisation *(inset*) from a holding potential *(V*_h_) of −80 mV in the absence (CTL; *left, black traces*) or presence of sevoflurane (1 MAC; 0.28 mM) (SEVO; *right, orange traces*). (*B,C*) Current-voltage relationship of channel activation displayed as normalized conductance (*G*/*G*_max_), and of inactivation (*I*_Na_/*I*_Na_max). Sevoflurane shifted the voltage dependence of activation by −6.2±3.2 mV (*V*_½-activ_−17.6±6.0 mV for CTL [*white circles*] *vs*. −23.7±6.0 mV for SEVO [*orange circles*], P=0.0026, n=7). Sevoflurane also shifted the voltage for half-maximal inactivation (*V*_½-inact_) by −6.1±2.7 mV toward hyperpolarized potentials (*V*_½-inact_ −44.8±7.0 mV for CTL [*white circles*] *vs*. −50.9±7.3 mV for SEVO [*orange circles*], P<0.0001, n=11). (*D*) This shift in *V*_½-inact_ was concentration-dependent, with the strongest effect of a −10.4±5.3 mV shift seen at 2 MAC sevoflurane. All data are mean±SD; **P<0.01, ****P<0.0001 *vs*. control by paired two-tailed Student *t*-test.

### Sevoflurane slows recovery from fast inactivation

We measured the effects of sevoflurane on recovery from inactivation using a holding potential of −120 mV, at which the majority of channels were in a resting, closed state (Fig. 3). Sevoflurane (1 MAC; 0.28 mM) significantly slowed the fast component of recovery, as reflected by an increase in the fast time constant τ_fast_, while the slow component τ_slow_ was unaffected (Supplementary Table S1). This indicates that sevoflurane selectively impairs the rapid recovery of Na_v_1.3 from fast inactivation, potentially prolonging the refractory period without altering slower recovery kinetics.

**Fig. 3.**
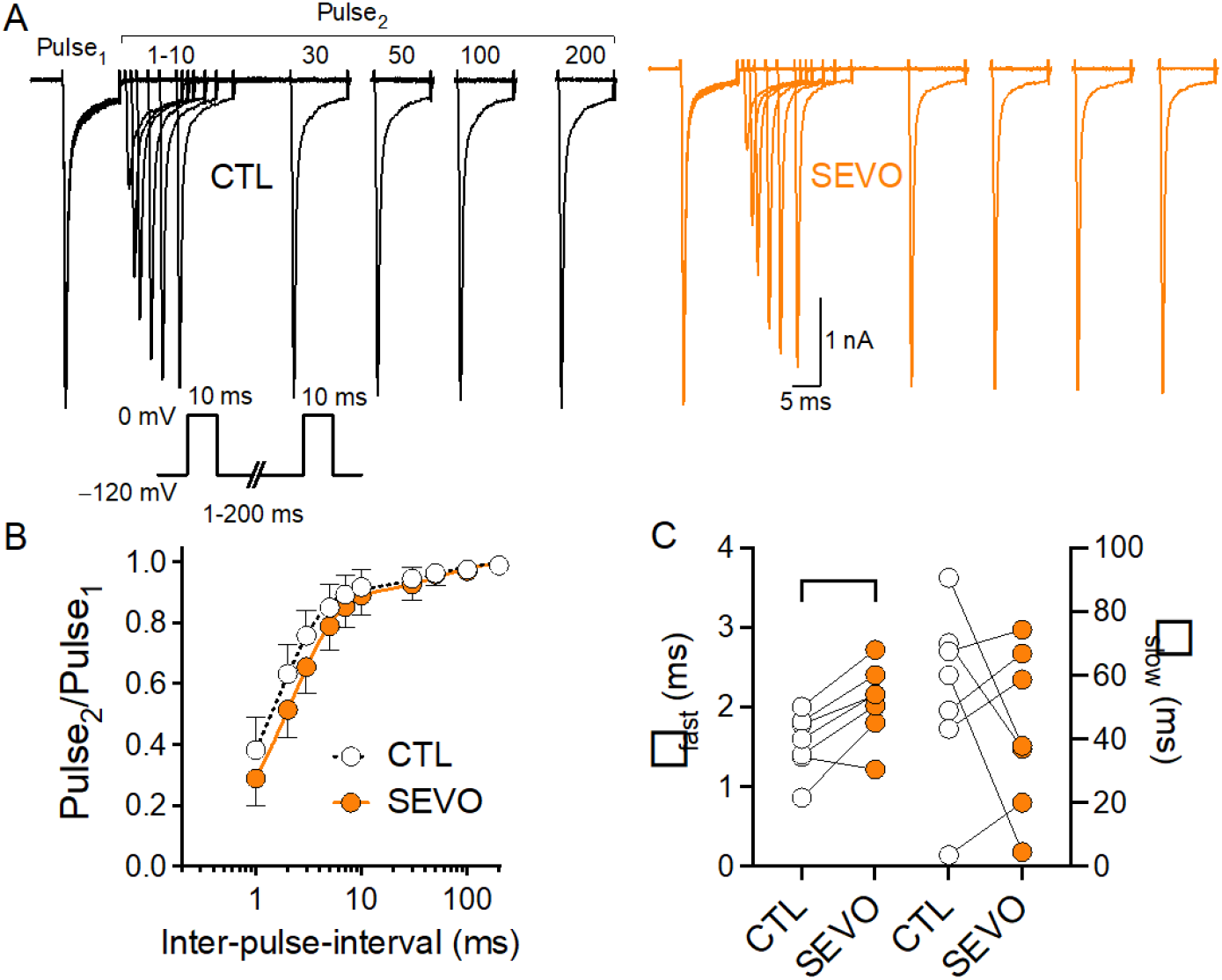
Effects of sevoflurane on Na_v_1.3 recovery from fast inactivation. We used a two-pulse protocol from a holding potential (*V*_h_) of −120 mV, with two 10 ms test pulses to 0 mV separated by an inter-pulse interval of 1–200 ms (*inset* in (A) shows stimulation protocol). (A) Overlaid macroscopic *I*_Na_ traces for the two pulses with increasing inter-pulse intervals in the absence (CTL, *black traces*) or presence (SEVO, *orange traces*) of sevoflurane (1 MAC; 0.28 mM). Peak *I*_Na_ of the second test pulse slowly recovers to control values with increasing inter-pulse durations. (B) Data fitted to a bi-exponential equation and plotted against inter-pulse-interval in the absence (*white circles, dotted line*) or presence (*orange circles, straight line*) of sevoflurane. (C) Recovery time constants τ_fast_ and τ_slow_ in the absence (*white circles*) or presence (*orange circles*) of sevoflurane. Sevoflurane increased τ_fast_ from 1.55±0.4 ms to 2.07±0.5 ms (P=0.0073), thereby slowing recovery from fast inactivation; τ_slow_ was not affected with 55±30 ms for CTL and 40±30 ms for SEVO; P=0.3789. Data are mean±SD, n=7, **P<0.01 *vs*. control by paired two-tailed Student *t*-test.

### Na_v_1.3 and Na_v_1.2 have distinct electrophysiological properties

We compared the effects of sevoflurane on Na_v_1.3 and Na_v_1.2 to investigate possible subtype-selective actions on developmentally distinct Na_v_ subtypes. Under control conditions, the kinetics of Nav1.3 were more depolarized compared to Na_v_1.2, with the voltage for half-maximal activation (*V*_½-activ_) and inactivation (*V*_½-inact_) significantly shifted by +10 mV (Fig. 2 and Fig. 4B,C, Supplemental Table S1,S2). These differences indicate that Na_v_1.3 channels exhibit depolarized gating, enhancing their likelihood of opening during action potentials and maintaining availability over a wider voltage range, and potentially influencing neuronal excitability.

**Fig. 4.**
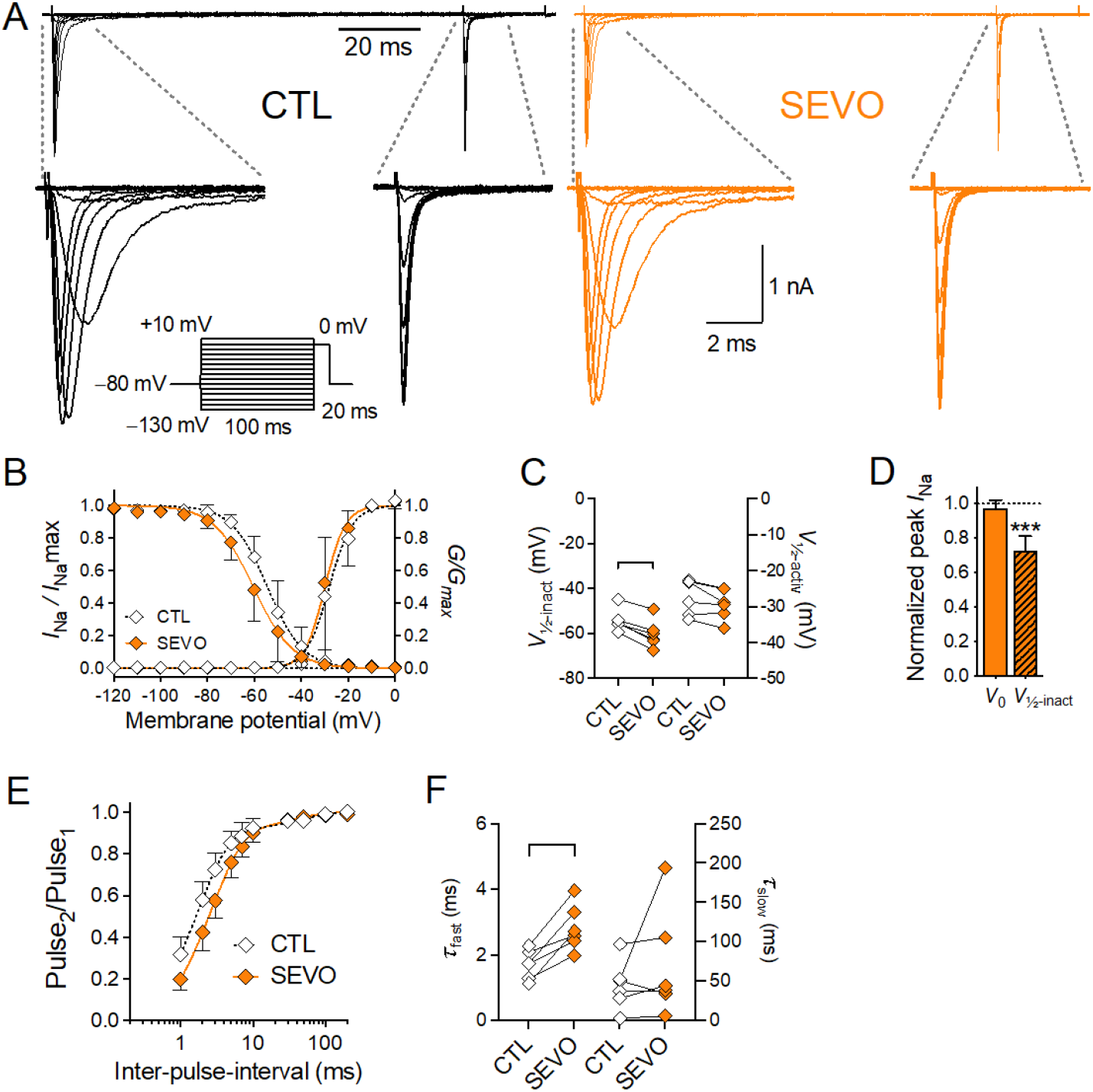
Effects of sevoflurane on Na_v_1.2 peak *I*_*Na*_, steady-state fast inactivation, and recovery from fast inactivation. (*A*) Representative families of whole-cell inward Na_+_ currents evoked by depolarization *(inset*) from a holding potential *(V*_h_) of −80 mV in the absence (CTL; *left*, black traces) or presence of sevoflurane (1 MAC; 0.28 mM) (SEVO; *right, orange traces*). (*B,C*) Current-voltage relationship of channel activation displayed as normalized conductance (*G*/*G*_max_) and inactivation (*I*_Na_/*I*_Na_max). Sevoflurane did not affect the voltage dependence of activation (*V*_½-activ_ −27.3±5.0 mV for CTL [*white diamonds*] *vs*. −29.4±4.2 mV for SEVO [*orange diamonds*], P=0.0544, n=6). However, sevoflurane significantly shifted the voltage for half-maximal inactivation (*V*_½-inact_) by −6.1±1.8 mV toward hyperpolarized potentials (*V*_½-inact_ −54.2±5.0 mV for CTL [*white diamonds*] *vs*. −60.3±6.2 mV for SEVO [*orange diamonds*], P<0.0005, n=6). (*D*) Effect of sevoflurane on peak *I*_Na_ (stimulation protocol see Fig 1A). Sevoflurane did not inhibit peak *I*_Na_ when the test pulse followed a prepulse to *V*_0_, a potential at which all channels were in the resting state. However, with a prepulse to *V*_½-inact_, a potential at which half of the channels were in the fast-inactivated state, sevoflurane produced a significant inhibition of peak *I*_Na_ (normalized peak *I*_Na_ 0.97±0.05 for *V*_0_, P=0.2238 and 0.73±0.08 for *V*_½-inact_, P=0.0005, n=6). (*E,F*) Recovery from fast inactivation was tested using the same protocol as in Fig 3.B. (*E*) Recovery data (Pulse_2_/Pulse_1_) was fitted to a bi-exponential equation and plotted against inter-pulse-interval in the absence (*white diamonds, dotted line*) or presence (*orange diamonds, straight line*) of sevoflurane. (*F*) Recovery time constants τ_fast_ and τ_slow_ in the absence (*white diamonds*) or presence (*orange diamonds*) of sevoflurane. Sevoflurane significantly increased τ_fast_ from 1.7±0.4 ms to 2.8±0.7 ms (P=0.0039), thereby slowing recovery from fast inactivation (τ_slow_ was not affected with 40±30 ms for CTL and 70.2±70 ms for SEVO; n=6, P=0.3257). See Supplemental Table S2 for all values. Data are mean±SD, **P<0.01, ***P<0.001 *vs*. control by paired two-tailed Student *t*-test.

Sevoflurane at a clinically relevant concentration of 1 MAC (0.28 mM) produced similar effects on Na_v_1.2 as observed with Na_v_1.3. Specifically, sevoflurane significantly inhibited peak *I*_Na_ of Na_v_1.2 when the test pulse followed a prepulse to *V*_½-inact_ (−56±7 mV), a potential at which ∼50% of channels are in a fast-inactivated state, but not with a prepulse to *V*_0_ (−120 mV) (Fig. 4D). The voltage dependence of steady-state fast inactivation was shifted by ∼ −6 mV toward more hyperpolarized potentials (Fig. 4B,C). Sevoflurane also slowed recovery from fast inactivation, with a significant increase in the fast time constant τ_fast_ (Fig. 4E,F). Sevoflurane did not significantly alter the activation kinetics of Na_v_1.2, highlighting a more pronounced effect on Na_v_1.3, where the drug modulates both activation and inactivation gating. This action might be particularly relevant in neurons where Na_v_1.3 is upregulated, such as during development or following injury.

### Effects of sevoflurane on persistent Na^+^ current

The distinct gating mechanisms of Na_v_1.3 result in *persistent Na*^*+*^ *current* (I_*Na*_*P*), which enhances repetitive firing and hyperexcitability during neuronal development ^34,36^. *I*_Na_P allows subthreshold depolarizations to generate APs, but it is unclear whether and how this is modulated by VAs. A large *I*_Na_P was generated by Na_v_1.3 which was significantly inhibited by 1 MAC sevoflurane (Fig. 5 A-C). *I*_Na_P generated by Na_v_1.3 was ∼ 6-fold greater than for Na_v_1.2, which was minimal and not further reduced by sevoflurane (–146±161 pA for Na_v_1.3 *vs*. –23±19 pA for Na_v_1.2; Fig. 5 D-F).

**Fig. 5.**
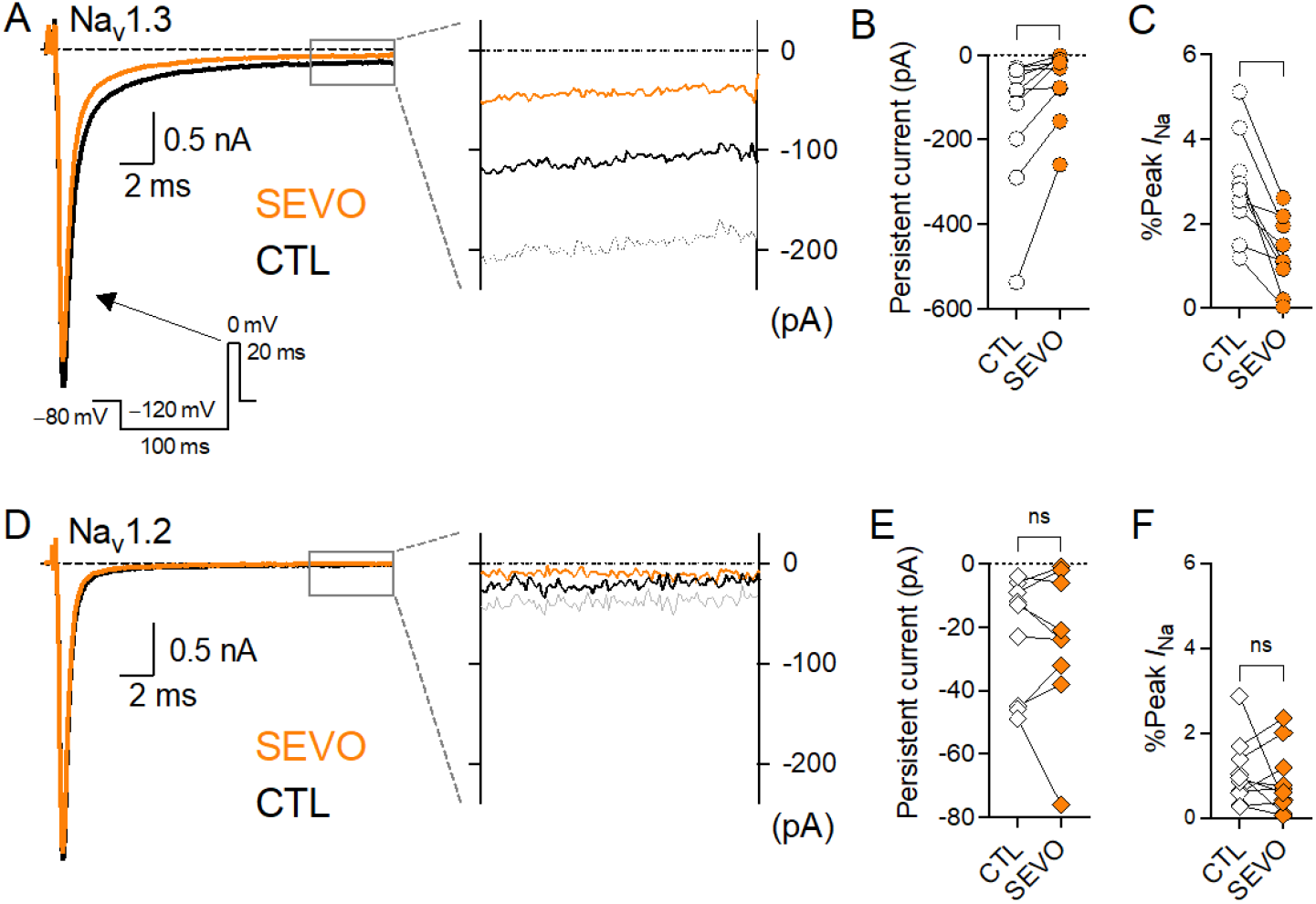
Effects of sevoflurane on Na_v_1.3 and Na_v_1.2 persistent Na^+^ current (*I*_Na_P). *I*_Na_P was tested using the depicted stimulation protocol (*A*, see *inset*). (*A,D*) Overlaid averaged macroscopic Na_+_ currents in the absence (*black traces*) or presence (*orange traces*) of sevoflurane (1 MAC; 0.28 mM). *Right panels* show a zoomed view of the final 5 ms of the 20 ms depolarization, highlighting *I*_Na_P (CTL *black traces*, SEVO *orange traces*, shaded areas represent SD). (*B, E*) Absolute *I*_Na_P values in pA, calculated as the mean current of the final 5 ms. Sevoflurane significantly reduced *I*_Na_P in Na_v_1.3 (from –146±161 pA for CTL [*white circles*] to –65±83 pA for SEVO [*orange circles*], P= 0.0144, n=10) but had no effect on Na_v_1.2 (from – 23±19 pA for CTL [*white diamonds*] to –25±23 pA for SEVO [*orange diamonds*], n=9). (*C, F*) *I*_Na_P expressed as a percentage of peak *I*_Na_, to account for variability in cell size and peak *I*_Na_ amplitude across recordings. Sevoflurane significantly reduced *I*_Na_P in Na_v_1.3 from 2.9±1.2% [CTL, *white circles*] to 1.3±0.9% [SEVO, *orange circles*], P<0.0011, n=9), but not in Na_v_1.2 (from 1.0±0.8% for CTL [*white diamonds*] to 0.7±0.6% for SEVO [*orange diamonds*], n=9). Data are mean±SD, *P<0.05, **P<0.01 *vs*. control by paired two-tailed Student *t*-test.

### Effects of sevoflurane on ramp currents

Inward Na^+^ currents during slow depolarizing ramps contribute to neuronal hyperexcitability by promoting repetitive firing. We examined the effects of sevoflurane on ramp-induced currents (*I*_Na_R) using a depolarizing voltage ramp from −120 mV to +40 mV over 800 ms (0.2 mV/ms). *I*_Na_R was quantified by integrating the inward *I*_Na_ over the voltage range from the activation threshold to the reversal potential, yielding the total charge transfer. To account for variability in channel expression levels among cells, this charge transfer was normalized to the peak *I*_Na_ elicited by a standard step depolarization, resulting in a normalized charge transfer value (pC/nA). Under control conditions, Na_v_1.3 mediated a greater normalized charge transfer during the ramp protocol compared to Na_v_1.2 (9.6±2.0 pC/nA for Nav1.3 *vs*. 3.5±1.0 pC/nA for Na_v_1.2; data from Fig. 6B). Sevoflurane at a clinically relevant concentration of 1 MAC (0.28 mM) significantly reduced the normalized charge transfer associated with *I*_Na_R in Na_v_1.3-expressing cells (Fig. 6A, B). In contrast, sevoflurane did not alter the normalized charge transfer in Na_v_1.2 (Fig. 6B), suggesting a subtype-specific effect on Na^+^ channel ramp currents.

**Fig. 6.**
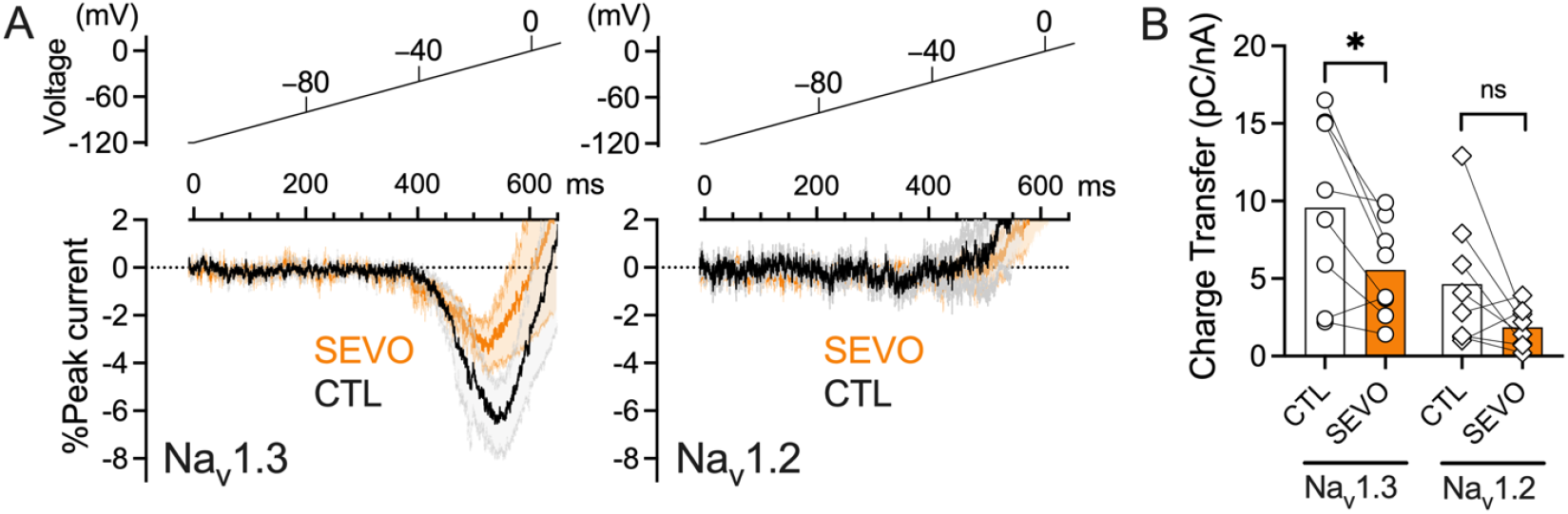
Sevoflurane selectively inhibits ramp-evoked Na^+^ currents (*I*_Na_R) in Na_v_1.3 but not Na_v_1.2. (*A*) Mean of all ramp currents recorded in response to a slow depolarizing ramp from –120 mV to +40 mV over 800 ms (0.2 mV/ms) for Na_å_1.3 (*left panel*) or Na_å_1.2 (*right panel*). Top panels show stimulation protocol. *Bottom panel* shows mean ramp current trace in the absence (CTL, *solid black trace*) or presence (SEVO, *solid orange trace*) of sevoflurane (1 MAC; 0.28 mM). The shaded areas show SEM for control (CTL, *grey area*) or sevoflurane (SEVO, *orange area*). Currents are normalized to peak *I*_Na_ and expressed as a percentage. (*B*) Normalized charge transfer (pC/nA) was calculated by integrating inward *I*_Na_ from the voltage at which currents activate to the voltage at which currents reverse polarity, followed by normalization to peak *I*_Na_. Under control conditions, Na_v_1.3 showed significantly greater normalized charge transfer than Na_v_1.2 (9.6±5.7 pC/nA for Na_v_1.3 *vs*. 3.5±2.7 pC/nA for Na_v_1.2, P=0.0226 by unpaired two-tailed Student *t*-test). Sevoflurane significantly reduced charge transfer in Na_v_1.3 (9.6±5.7 to 5.6±3.1 pC/nA, P=0.0203, n=8), but not in Na_v_1.2 (3.5±2.7 to 1.7±1.3 pC/nA, P=0.1680 n=7). Data are presented as mean±SD. *P<0.05 *vs*. control by paired two-tailed Student *t*-test.

## Discussion

The developmentally expressed sodium channel Na_v_1.3 is sensitive to modulation by the volatile anesthetic sevoflurane. While the effects of volatile anesthetics on Na_v_1.2 have been characterized previously, their impact on developmentally expressed Na_v_1.3 were previously unknown. We compared the effects of sevoflurane on the channel gating properties of human Na_v_1.3 and Na_v_1.2, the predominant Na^+^ channel subtypes expressed in immature and mature neurons, respectively. Under control conditions, Na_v_1.3 exhibited gating at more depolarized potentials compared to Na_v_1.2: both the voltage of half-maximal activation (*V*_½-activ_) and inactivation (*V*_½-inact_) were ∼10 mV more positive in Na_v_1.3. Despite these differences, sevoflurane produced a comparable hyperpolarizing shift in steady-state fast inactivation for both subtypes. The degree of voltage-dependent inhibition of peak *I*_Na_ and the slowing of recovery from fast inactivation were also similar between Na_v_1.3 and Na_v_1.2, suggesting that baseline differences in gating do not substantially alter the overall pattern of modulation by sevoflurane.

Persistent Na^+^ current (*I*_Na_P) can drive intrinsic neuronal hyperexcitability^47^ and has been implicated in various physiological and pathological processes, including pacemaking, neuronal injury, chronic pain, and epilepsy. We observed large *I*_Na_P in cells expressing Na_v_1.3, consistent with enhanced excitability of immature neurons. Sevoflurane reduced *I*_Na_P in Na_v_1.3 expressing cells, while Na_v_1.2 had much smaller persistent currents than Na_v_1.3, and were not further reduced by sevoflurane.

Ramp currents (*I*_Na_R), which can enhance responses to subthreshold depolarizations by reducing action potential firing threshold^48^, were more pronounced in Na_v_1.3 expressing cells. This observation is consistent with studies indicating that mutations in Na_v_1.3 linked to neuronal hyperexcitability increase ramp and persistent currents. Selective inhibition of these currents could mediate developmental stage-specific effects of sevoflurane and possibly other anesthetics on neuronal excitability.

Multiple volatile anesthetics have been shown to inhibit all neuronal Na_v_ subtypes, including Na_v_1.1, Na_v_1.2, and Na_v_1.6, leading to reduced AP amplitude in mature neurons at clinically relevant concentrations^11,15,31,33^. In addition to shifting the voltage dependence of steady-state fast inactivation and slowing channel recovery, our findings show that sevoflurane also reduces *I*_Na_P and *I*_Na_R in Na_v_1.3-expressing cells, providing a basis for previous observations that volatile anesthetics suppress transient Na^+^ current (*I*_Na_T) and *I*_Na_P, thereby reducing hippocampal excitability^40^. Thus, during brain maturation, multiple Na_v_1.3 gating mechanisms are likely targeted by volatile anesthetics to inhibit Na_v_-activated Ca^2+-^influx and neurotransmitter release that contribute to the neurophysiological actions of anesthetics.

Modulation by anesthetics of neuronal activity during early development during a sensitive developmental window can lead to persistent alterations in neuronal function. The neurotoxic effects of general anesthetics administered at critical periods early in development in the mammalian brain ^51-53^ are associated with structural and functional changes, including alterations in dendritic spine dynamics, dendritic arborization, and synaptic protein expression^51-53^. Since general anesthetics have selective effects on neurotransmission by depressing excitatory and enhancing inhibitory synaptic transmission, the balance of excitotoxicity from glutamatergic transmission in immature neuronal networks can also affected by early exposure to anesthetics^54,55^. Alterations in neuronal excitability by general anesthetics with cell type specificity have been observed in cortical, thalamocortical, and hippocampal networks^56-59^. In the motor cortex, neonatal propofol anesthesia reduces neuronal activity in excitatory and inhibitory neurons, which is associated with motor learning impairments. In contrast, administration of agents that attenuate GABA_A_ receptor activity or potentiate AMPA signaling during emergence from anesthesia can ameliorate both neuronal circuit dysfunction and learning deficits^60^. In contrast, in the somatosensory cortex of young mice, sevoflurane increases neuronal activation and behavioral hyperactivity, while propofol attenuates this neuronal activation via enhancement of GABA signaling^61^. As Na_v_1.3 is predominantly expressed in excitatory neurons^62^ and drives persistent excitability, our findings suggest a major role for Na_v_1.3 in cell type-specific and agent-specific effects of anesthetic exposure to enhance excitability of immature brain networks, warranting further investigation.

Mature neuronal circuits are also susceptible to functional changes following anesthetic exposure. Perioperative neurocognitive disorders (PND), including delirium, cognitive dysfunction, and delayed cognitive recovery, can occur following anesthesia and surgery^63-65^. Older patients are particularly affected, but additional risk factors include pre-existing cognitive impairment or disease^63,66^. Following injury or disease, excitatory neurons augment their high-frequency firing by upregulating Na_v_1.3 expression^37^. Significant increases in Na_v_1.3 mRNA and protein levels in the spinal cord and thalamus^67-70^ are found in a range of neuropathic pain models and conditions, including dorsal root ganglion axotomy^37,71,72^, spinal nerve ligation^73,74^, chronic constriction injury (CCI^71,75,76^, spared nerve injury (SNI^62,77^, trigeminal neuralgia^78^, diabetic neuropathy^79^, and post-infection neuralgia^80^. Activity of Na_v_1.3 is associated with spontaneous ectopic firing and development of neuropathic pain as selective inhibition of Na_v_1.3 function by gene ablation, antisense knockdown, or viral-mediated knockdown reduces allodynia and hyperalgesia associated with SNI, CCI, and diabetic neuropathy^75,81-83^. Increased Na_v_1.3 expression following CNS injury might interact with anesthetic exposure to modulate overall neuronal excitability at the network level in these conditions.

Our baseline Na_v_1.3 activation properties (*V*_½-activ_) including *I*_Na_P are consistent with previous reports^49,50^. However, our *V*_½-inact_ values were notably more depolarized at ∼–45 mV compared to ∼–70 mV in those studies. Additionally, we observed larger ramp currents and normalized charge transfer in Na_v_1.3-expressing cells than those reported by Vanoye et al.^49^. These differences might stem from alterations in holding potentials (*V*_h_) (we chose a more physiological *V*_h_ for certain experiments) or differences in auxiliary subunit expression, as we co-expressed only the β1- subunit, whereas Vanoye et al. included both β1 and β2. The β2 subunit is known to modulate inactivation gating and reduce persistent current amplitude^84-86^, which might account for the more depolarized inactivation and enhanced ramp currents observed in our study.

Multiple binding sites have been proposed for volatile anesthetic interactions with Na_v_ ^87-89^; further studies are required to determine whether differences in anesthetic binding to specific sites contribute to these subtype-selective effects of sevoflurane. Various volatile anesthetics might exhibit different affinities for Na_v_ subtypes^90,91^, with sevoflurane showing stronger interactions with Na_v_1.3 compared with Na_v_1.2. Moreover, Na_v_ subtypes not only show cell-type specific expression^7^ but also distinct subcellular and cellular localizations^8^. Parvalbumin-expressing interneurons are enriched in Na_v_1.1, while excitatory pyramidal neurons express more Na_v_1.2, Na_v_1.3, and Na_v_1.6^7,62,92,93^. While all Na_v_ subtypes are found at the axon initial segment (AIS) to some degree, Na_v_1.1 and Na_v_1.2 are preferentially expressed in the proximal AIS^94^, potentially allowing direct targeting of AP initiation by anesthetics.

Further studies of the selective effects of sevoflurane on Na_v_ subtypes could lead to the development of more selective and potentially safer anesthetic agents that selectively target specific Na_v_ subtypes based on neuronal stage or neuropathological state by reducing deleterious off target effects or specifically targeting desired effects.

## Abbreviations

Na_v_: voltage-gated sodium channel
SEVO: sevoflurane
VA: volatile anesthetic
AP: action potential
TEA-Cl: tetraethylammonium chloride

## Acknowledgments

We thank Prof. Jennifer Kearney (Northwestern University) for generously providing the human Na_v_1.3 clone and for her valuable assistance with expression of Na_v_1.3 in our cell lines. We also thank Nicolas Lewis for his help with plasmid amplification.

## Funding

US National Institutes of Health grant R01 GM58055 to HCH, and R01 GM130722 to JP.

## Supplementary Information for

**Table S1.**
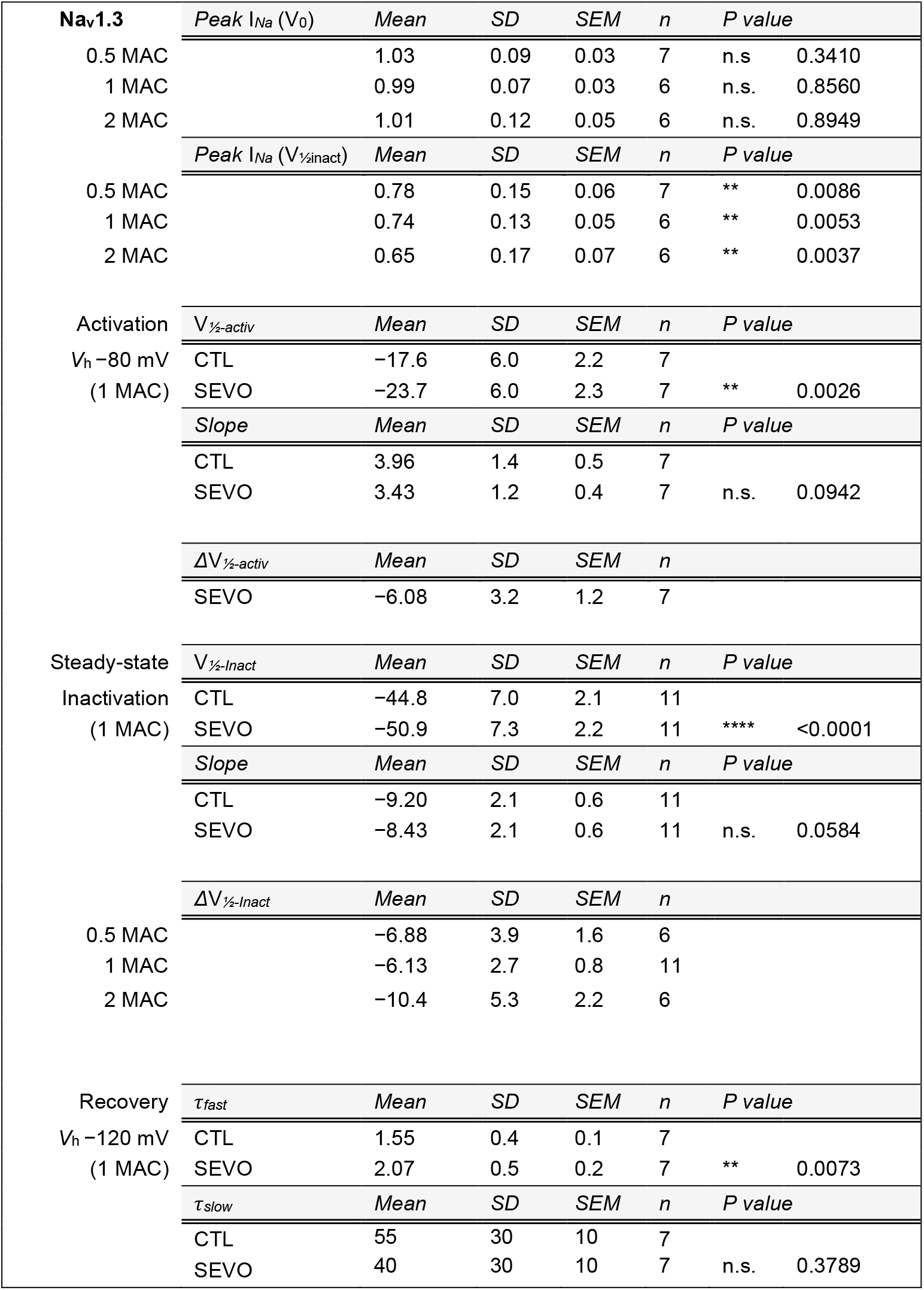
Effects of sevoflurane on Na_v_1.3

**Table S2.**
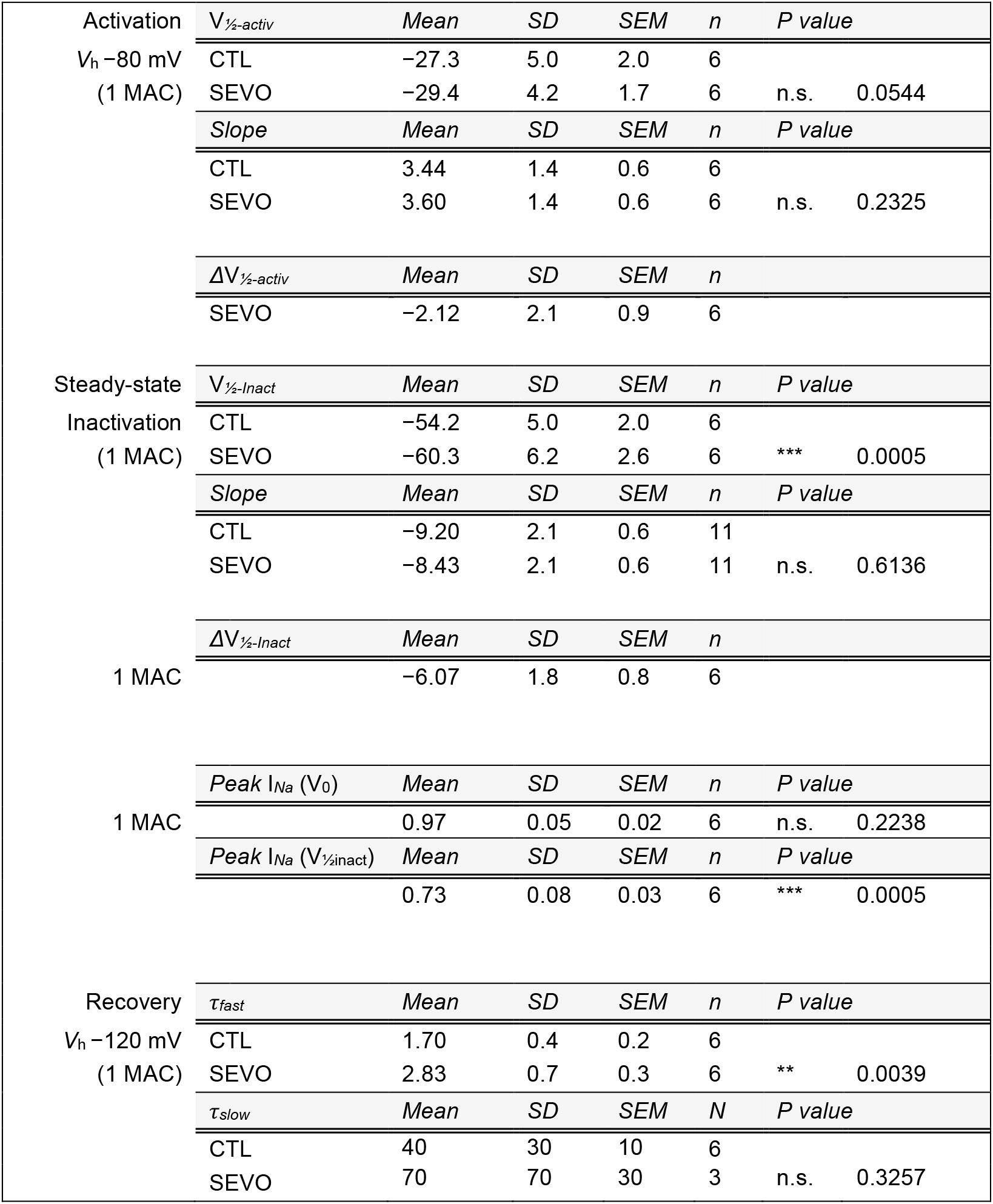
Effects of sevoflurane on Na_v_1.2

## Notes

### Competing Interest Statement

The authors have declared no competing interest.

